# OddSNP: a predictive framework for optimizing multiplexed single-cell RNA-seq experiments

**DOI:** 10.64898/2025.12.08.692882

**Authors:** Rodolfo S. Allendes Osorio, Toshiya Nishimura, Yuichi Shigihara, Masaki Kimura, Takanori Takebe, Takahiro Nemoto

**Affiliations:** Premium Research Institute for Human Metaverse Medicine (WPI-PRIMe), The University of Osaka, 2-2 Yamadaoka, Suita-shi 565-9315, Osaka, Japan; Center for Stem Cell and Organoid Medicine (CuSTOM), Division of Gastroenterology, Hepatology and Nutrition, Division of Developmental Biology, Cincinnati Children’s Hospital Medical Center, 3333 Burnet Avenue, Cincinnati, 45229, OH, USA

**Keywords:** scRNAseq, demultiplexing, SNP, human liver organoids

## Abstract

Donor multiplexing is a powerful strategy to increase scale, lower the costs, and reduce batch effects in single-cell RNA sequencing (scRNAseq), but clear guidelines for experimental design are lacking, forcing researchers to risk costly demultiplexing failures. To address this, we introduce SNP-Information Content (SNP-IC), a quantitative metric computable from simple unpooled pilot data that accurately predicts the success of genotype-based demultiplexing. Across multiple large-scale datasets using stem cell and organoid models, we establish a robust SNP-IC threshold of approximately 50, above which cells can be reliably assigned to their donor of origin. For more challenging genotype-free approaches, we define a pairwise metric, cpSNP-IC, and demonstrate a much higher requirement of approximately 3,000. Our open-source framework, *oddSNP*, implements this predictive model, allowing researchers to perform in silico titrations of sequencing depth and donor complexity to optimize experimental design before committing to large-scale studies. *oddSNP* provides a practical framework, enabling researchers to strategically optimize sequencing depth and donor numbers to maximize experimental success while managing costs and minimizing the risk of catastrophic data loss.

## 1 Introduction

Rapid advances in single-cell RNA sequencing (scRNAseq) have revolutionized our ability to explore transcriptomic landscapes at single-cell resolution [1, 2, 3, 4, 5, 6, 7]. However, the use of scRNAseq in multi-donor large population cohorts remains limited, mainly due to sample availability, technical batch effects, and cost. To address these challenges, the generation of samples pooled from different donors that can be processed together in a single scRNAseq experiment (multiplexing) has emerged. The key step in multiplexing experiments is *de-multiplexing*, where the origin of the donors is traced back from each cell data. Two distinct strategies exist for this purpose [8]. The first involves the use of add-on barcodes (“hashing”). In this category, DNA-tagged antibodies targeting ubiquitous surface proteins (Cell Hashing [9]) or lipid-anchored oligos (MULTI-seq [10]) are added to label samples prior to pooling. The barcode is co-captured and counted to assign sample identity and detect inter-sample doublets. These methods are broadly applicable across species, work for both cells and nuclei, and are supported by commercial kits (e.g., 10x CellPlex). However, they introduce additional wet-lab steps and require extra sequencing reads for tag counting. In contrast, the second strategy works without adding any DNA barcodes. Instead, it infers each cell’s donor of origin directly from cDNA reads overlapping polymorphic sites, significantly simplifying the workflow while still enabling multiplexing experiments (Fig.1a).

**Figure 1:**
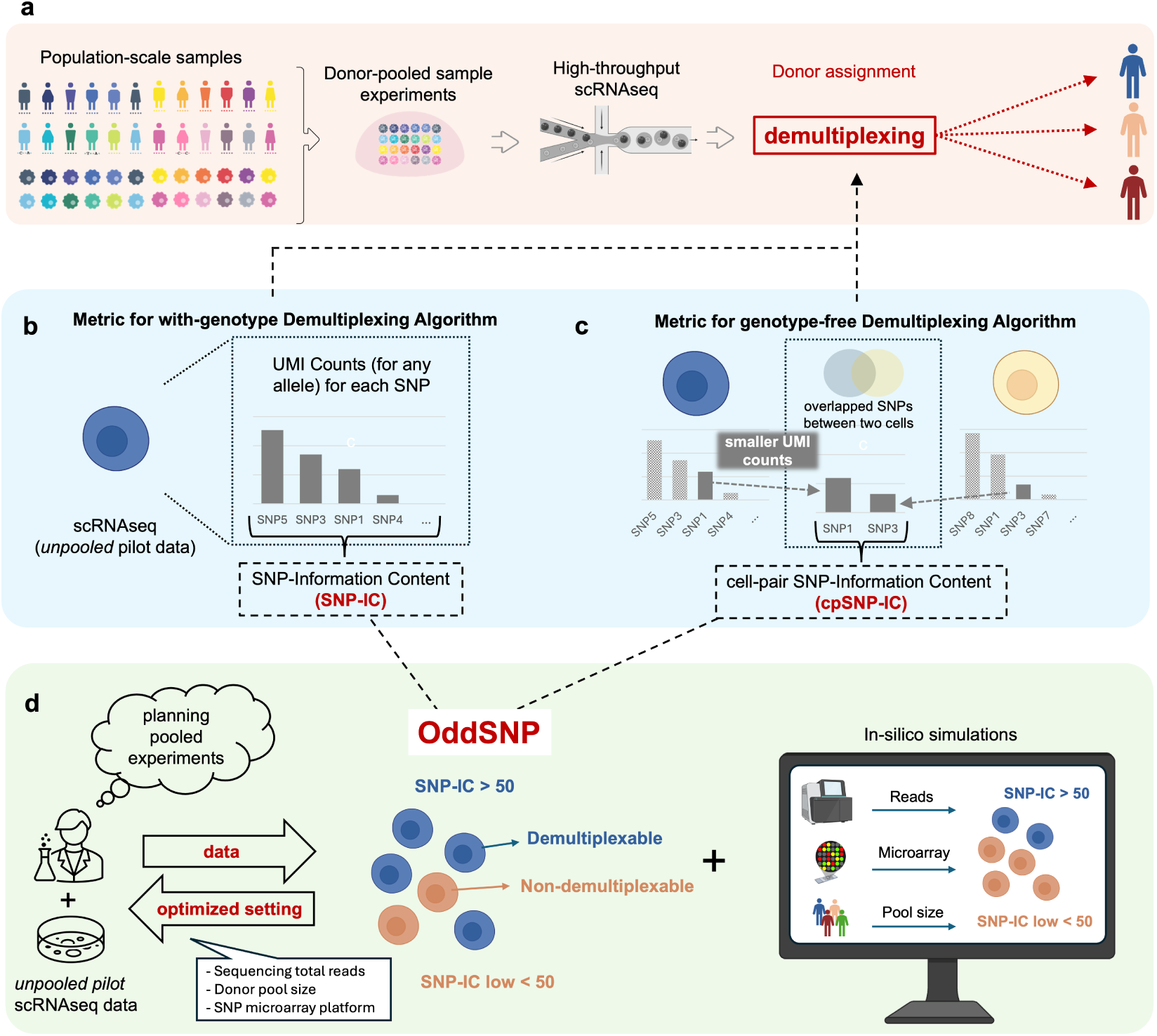
Overview of oddSNP. (a) Samples from multiple donors are pooled together, and scRNAseq is performed to obtain transcriptional profiles at the single-cell level. The goal of demultiplexing is to assign each cell to its donor of origin. (b) To quantify the ability of each cell to be demultiplexed in with-genotype algorithms (See Fig.S1), we introduce SNP-IC, defined as the sum of UMI counts across identified SNPs within the scRNA seq data. (c) Similarly, to quantify the ability of a pair of cells to be distinguished in genotype-free algorithms (See Fig.S1), we introduce cpSNP-IC, defined as the sum of UMI counts over shared SNPs between the two cells. (d) Both SNP-IC and cpSNP-IC can be computed from unpooled scRNAseq data using our proposed pipeline, *oddSNP*. This enables users to estimate, for future pooled experiments, the expected proportion of cells that can be correctly demultiplexed using with-genotype algorithm (based on SNP-IC), as well as the proportion of cell pairs that can be distinguished using genotype-free algorithms (based on cpSNP-IC). oddSNP also includes a downsampling function, enabling users to explore optimal experimental conditions *in-silico*.

These polymorphism-based methods are expanding their applicability. For example, Perez et al. [11] used the sequencing results from a pool of 264 samples to study molecular and genetic associations to lupus. Both Neaving et al. [12] and Wells et al. [13] use the concept of *village* to describe experimental platforms where cells from different donors are pooled and cultured together. Furthermore, the same approach was also employed to study cancer cell lines by Weber *et al.* [14]. The benefits of using pooled samples also apply to the generation of organoids [15], simple 3D cellular structures cultured in vitro used to simulate the corresponding tissue in vivo. For example, Kimura et al. [16] developed and used a pooled human population organoid panel (PoP) to study the influence that specific Single Nucleotide Polymorphisms (SNPs) have on the onset of Metabolic dysfunction associated steatohepatitis (MASH). Caporale *et al.* also used the same strategy to study cortical brain organoids for the longitudinal dissection of developmental traits [17]. Again, the use of a PoP has the combined advantage of avoiding batch effects that could arise from single-donor generated organoids, together with the potential to study the impact that genotype variants have on phenotype without the complexity added by external factors such as the donor’s metabolic status.

The polymorphism-based pooled strategy has proven to be useful and has been applied in many settings. However, to the best of our knowledge, systematic guidelines for designing experimental setups and selecting appropriate parameters are still lacking. Such guidelines are particularly important when sufficient coverage of exonic variants, required to identify donors within pooled samples, may not be available. In practice, these guidelines depend on the types of demultiplexing algorithms used: those that require independent genotyping of donors included in the pooled dataset, such as Demuxlet [18], Dropulation [13], Vireo (with-genotype) [19] and Demuxalot [20] (hereafter referred to as with-genotype algorithms, shown in Fig.S1-left); and those that do not, such as Freemuxlet [21], Souporcell [22], Vireo (genotype-free) [19] and ScSplit [23] (referred to as genotype-free algorithms, shown in Fig. S1-right). In this article, we address this gap by providing an in-silico platform to identify optimal experimental conditions, enabling users to confidently assess whether polymorphism-based demultiplexing is suitable before conducting pooled experiments.

To this end, we systematically analyze pooled data from a large set of 104 distinct samples [12], 24-donor liver organoid pooled data [16], and our novel in-house dataset comprising mixtures of 200 samples. Specifically, we introduce SNP-Information Content (SNP-IC; Fig.1b) and pair-cell SNP-IC (pcSNP-IC; Fig.1c) as measures for assessing whether each cell can be properly demultiplexed. Across multiple datasets, we observed that SNP-IC≃ 50 serves as the threshold at which with-genotype algorithms can reliably demultiplex cells, while pcSNP-IC≃ 3, 000 is required for genotype-free algorithms. We propose a framework, *oddSNP* (Optimizing demultiplexing pooled experiments based on Single-Nucleotide Polymorphism), that computes these measures for each cell barcode from *unpooled* scRNAseq datasets. Within oddSNP, users can also perform in-silico simulations under various conditions, such as varying reads per cell and numbers of SNPs. This enables systematic evaluation of demultiplexing feasibility across diverse scenarios (Fig.1d). The oddSNP framework is available at [URL to be inserted], freely accessible to anyone planning to perform pooled experiments.

## 2 Results

### 2.1 Comparison of eight demultiplexing algorithms

Currently, several algorithms are available for the demultiplexing of pooled scRNAseq datasets. When applied to the same dataset, they may produce different classification results in principle, thus effective strategies for their evaluation are critical, for discussing demultiplex-ability criterions. Building on the consensus approach described in [17] and leveraging the Demuxafy platform [24], we benchmark eight algorithms: four with-genotype (Dropulation, Demuxlet, Demuxalot, and Vireo-Informed) and four genotype-free (Freemuxlet, Vireo-Uninformed, Souporcell, ScSplit). We apply these algorithms to demultiplex the 104-donor cell village dataset [13], consisting of more than 3.6 million cells, from 104 original donors, pooled and cultured together in equal proportions (See Methods for more details). In Fig. 2, a comparison of the demultiplexing results obtained using different algorithms (panels a and c), along with a summary of the running time required for each algorithm (panel b), is presented. With the exception of Demuxlet and Freemuxlet, all the methods tested were able to complete execution (See Methods for more details).

**Figure 2:**
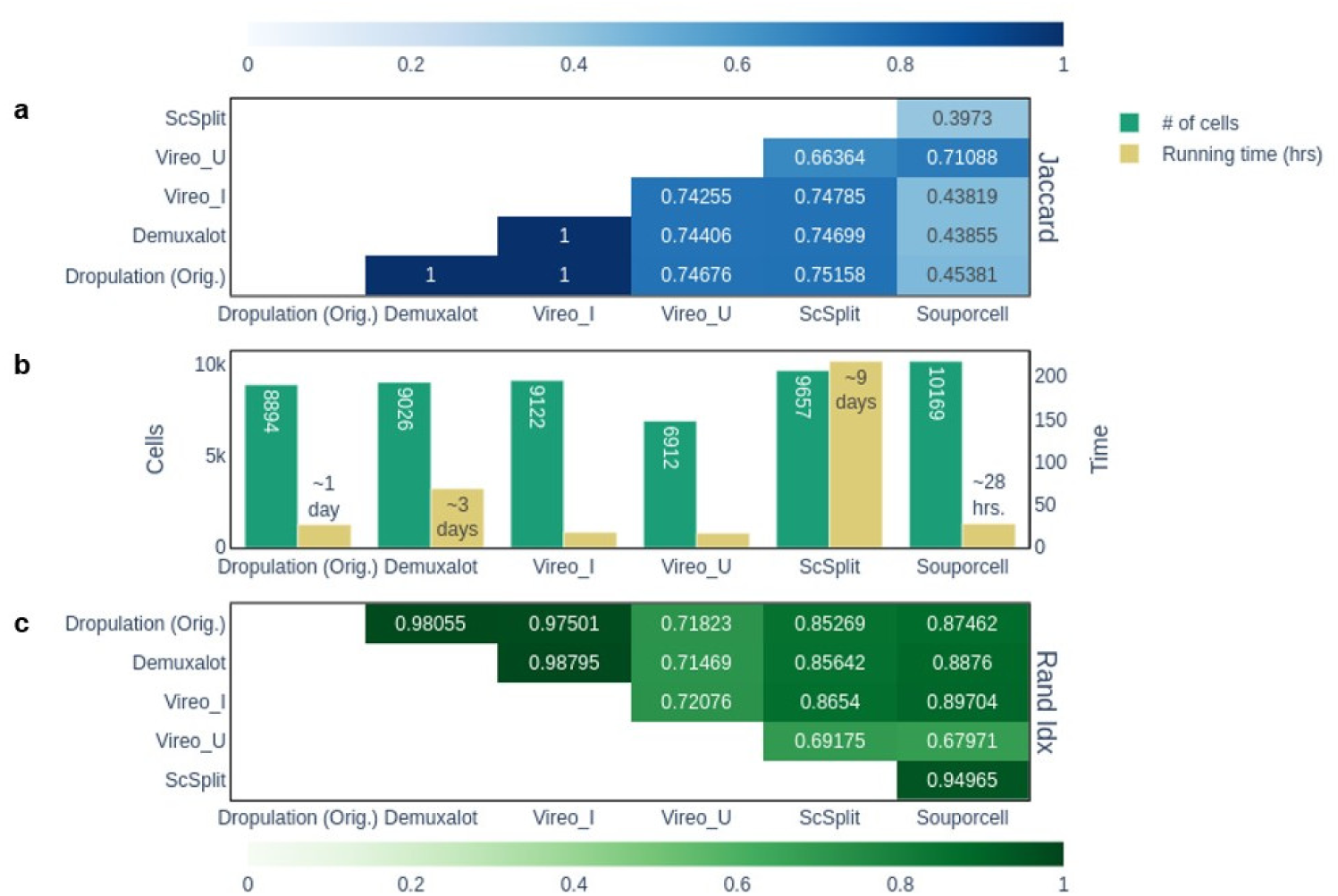
Pairwise comparison of various demultiplexing algorithms on the 104-donor cell village dataset. (a) The overlap between cells classified as singlets by both methods is estimated using the Jaccard Index. Higher values indicate greater overlap. (b) The number of cells that are mapped to a single donor (*i.e.*, singlet cells) is shown in green, while the approximate wall time required for execution is shown in yellow. (c) The agreement in the actual donor identity of singlet-classified cells between the two methods is evaluated using the Rand Index. Higher values indicate stronger agreement. Demuxlet and Freemuxlet runs are excluded due to technical reasons (Methods for details).

To assess the similarity between methods in a pairwise manner, we use two measures: the Jaccard index[25] and Rand index[26] (see Methods sections 4.4 and 4.7). The Jaccard index [25] evaluates the overlap between the two sets of cells that each method identifies as singlets, while the Rand index assesses the agreement in the actual identity of the cells classified in both methods as singlets. Both measures take values ranging from 0 to 1, with values closer to 1 indicating stronger agreement between the two methods. Fig.2 suggests that with-genotype algorithms (Vireo-I, Dropulation, Demuxalot) yield highly consistent results, both in overlap (Jaccard index = 1) and in cell identity (Rand index *>* 0.97). This strong agreement across algorithms indicates that their results are robust and reliable.

On the other hand, genotype-free algorithms produce results with small inconsistencies across methods. Vireo-U shows Jaccard and Rand indices of about 0.7-0.75 compared with other algorithms, except for Souporcell. Souporcell achieves a high Rand index with with-genotype algorithms (0.85-1), but its Jaccard indices against other algorithms are much lower (∼0.45). Among the genotype-free approaches, ScSplit performs best, reaching a high Rand index with genotype-based algorithms (0.85-1) and moderate Jaccard indices (∼0.75). However, it is worth noting that ScSplit requires about 10 times more computation time than Vireo-U and Souporcell (Fig. 2b).

Overall, with-genotype algorithms consistently demonstrate similar performance across different methods, whereas genotype-free algorithms exhibit more heterogeneous performance in terms of both accuracy and computational time. As a representative algorithm, we select Vireo for the criterion analysis below, since it is available in both with-genotype and genotype-free versions (*Vireo-I* and *Vireo-U*, respectively) and is among the fastest methods. Among with-genotype algorithms, we expect that other methods to show comparable criteria, while among genotype-free algorithms, Vireo achieves intermediate accuracy, making it a suitable representative for analyzing the criterion.

### 2.2 Demultiplex-ability analysis for with-genotype algorithms

When de-multiplexing cells, the number of SNPs identified from each cell’s mRNA-derived sequences would strongly influence demultiplexing ability. In with-genotype algorithms, the core idea (although the details should be different) is to compare cell-specific SNPs against reference SNP panels from the donors for assigning probabilities of donor origin. Consequently, the amount of SNP information, e.g., the number of detected SNPs and sequencing depth, is the primary factor affecting the demultiplexing ability. By contrast, genotype-free algorithms operate without donor reference genotypes and instead rely on direct comparisons of SNP profiles across cells to identify groups originating from the same donor. This framework indicates that, in addition to the amount of SNP information, the composition of the donor mixture, e.g., diversity among donors or how many donors are mixed together inside the same pool, also plays a critical role in demultiplexing accuracy (See Supplementary Fig.S2 for more details).

Based on this observation, for assessing demultiplexing performance of with-genotype algorithms, we introduce the SNP information content (SNP-IC), as the sum of UMI counts over the identified SNPs in that cell (Fig.1b). To systematically investigate how this quality affects the demultiplexing performance, we generated a series of down-sampled datasets from the original scRNAseq data by randomly selecting 50%, 10%, 1%, 0.5% and 0.1% of the original reads. First of all, we observed that even when the number of reads is significantly reduced (e.g. to 0.1%), the total number of cells in each filtered selection remains nearly the same, as indicated by the bar heights in Fig. 3a). To evaluate the demultiplexing outcomes for each down-sampled dataset, we used the demultiplexing results from the original dataset (100%) as the ground truth. We defined true positives as cells classified as singlets whose assignments matched those in the original data (100%), and false positives as cells classified as singlets whose assignments differed from the original data. The outcomes were clearly affected by downsampling, as shown in the corresponding bar graph. Whilst at lower number of reads, most or all cells are unassigned (*i.e.*, no reliable information can be inferred from the algorithm), once around 10% of the original reads become available, the cell mappings become stable. The assignment of donors to each cell is also consistent at sequencing depths of ≥ 10%, as illustrated by the high mapping accuracy (Fig. 3a).

**Figure 3:**
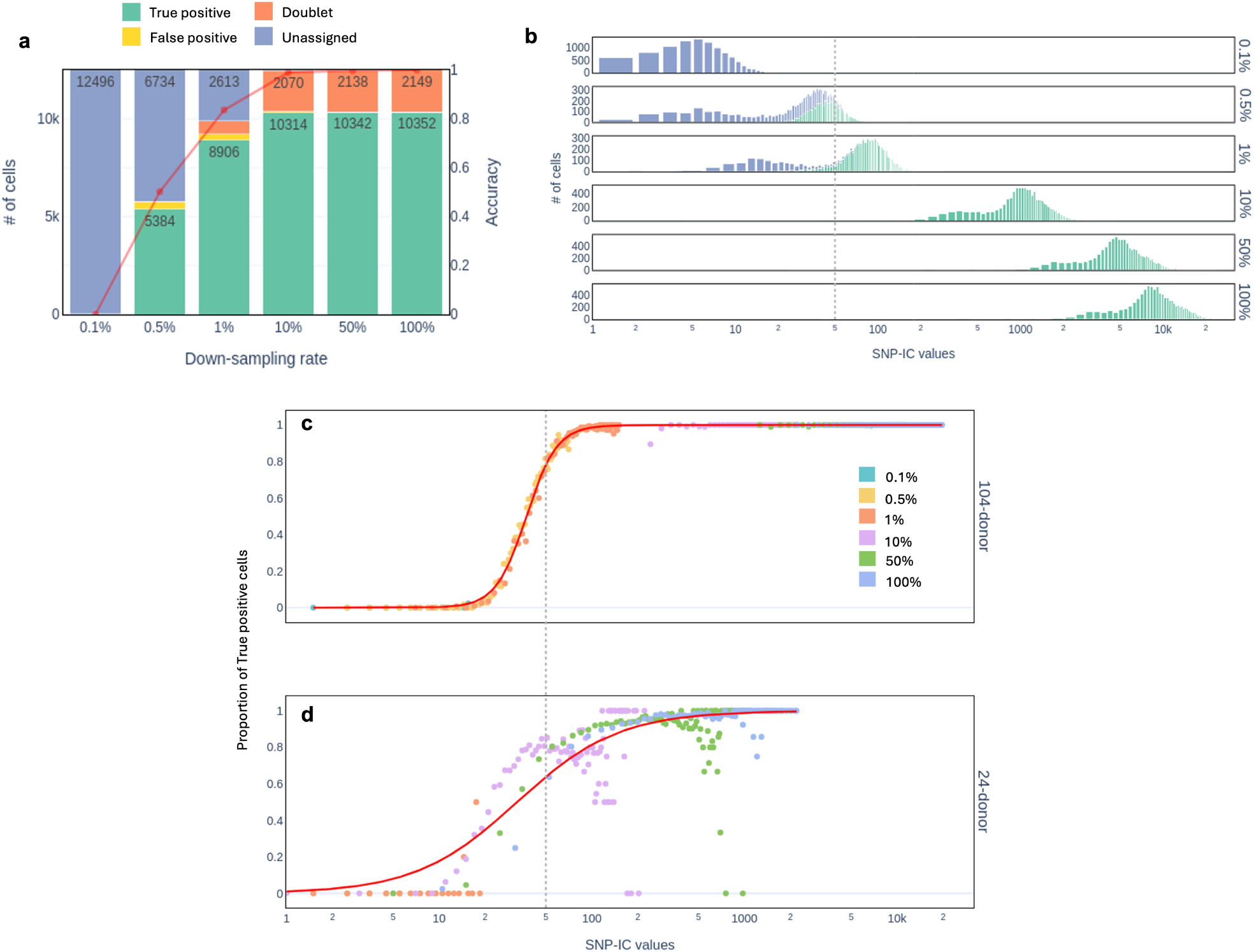
Demultiplex-ability analysis for with-genotype algorithms. (a) With-genotype Vireo classification results (singlet-false-positive, singlet-true-positive, doublets, unassigned) in down-sampled datasets, where downsampling is performed based on sequencing depth (0.1% - 100%) using the 104-donor cell village dataset. The demultiplexing accuracy is shown as a red solid line. (b) Histogram of SNP-IC values for each down-sampled level (top - 0.1%; bottom - 100% of reads). Different classification categories are shown in distinct colors. The dashed line indicates SNP-IC=50, which can serve as a threshold for correctly demultiplexed cells. In this panel, only cells categorized as singlets in the original data (100%) are shown. (c) Proportion of true-positive cells as a function of SNP-IC for each bin used in panel (b). (d) The same proportion for 24-donor HLO dataset. Note that in panels (c) and (d), the set of SNPs used to compute SNP-IC matches the set used to run the with-genotype Vireo. Panels (c) and (d) demonstrates that the SNP-IC threshold is robust across different datasets, as well as to variations in the SNP set used for computing SNP-IC.

To investigate the relationship between demultiplexing performance and SNP-IC, in Fig.3b we present the histogram of this quantity for each cell, stratified by the demultiplexing outcomes. As can be seen, when the SNP-IC increases, the number of unassigned and false positive cells decreases. In particular, there appears to be a threshold around

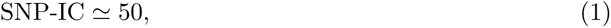

that determined whether a cell can be successfully demultiplexed. This effect is clearly illustrated in Fig.3b, where the proportion of true positive cells is plotted against the SNP-IC.

To demonstrate the generality of our SNP-IC threshold criterion, we performed the same downsampling analysis using a different pooled scRNAseq dataset of induced pluripotent stem cell (iPSC)-derived human liver organoids (HLO), consisting of 24 donors [16]. As shown in Fig.3d (and histogram in Supplementary Fig.S4b), even though the data come from different cell types, involve different numbers of donor mixtures, and represent distinct genotype backgrounds, we observed consistent results for the SNP-IC threshold around 50, suggesting the robustness of our criterion. Furthermore, note that in Fig.3c and 3d, the set of SNPs used to compute SNP-IC matches the set used to run the with-genotype Vireo. Specifically, the SNP list from the OmniExpressExome array was used for (c), while a more complete list derived from Whole-genome sequencing was used for (d)). This further demonstrates that the SNP-IC threshold is robust to variations in the SNP set used to compute SNP-IC.

### 2.3 Demultiplex-ability analysis for genotype-free algorithms

Similarly, we also evaluated the SNP-IC when using genotype-free algorithms. Using the same down-sampled datasets described above, we ran the genotype-free version of Vireo for demultiplexing. As shown in Fig.4a (and Supplementary Fig.S3b), we observe that the number of singlets does not increase monotonically with sequencing depth. This occurs because, at lower read depths, more cells categorized as singlets are incorrectly assigned compared to those at higher read depths. Consistent with this, the mapping accuracy increases monotonically with depth (the red solid line in Fig.4a).

**Figure 4:**
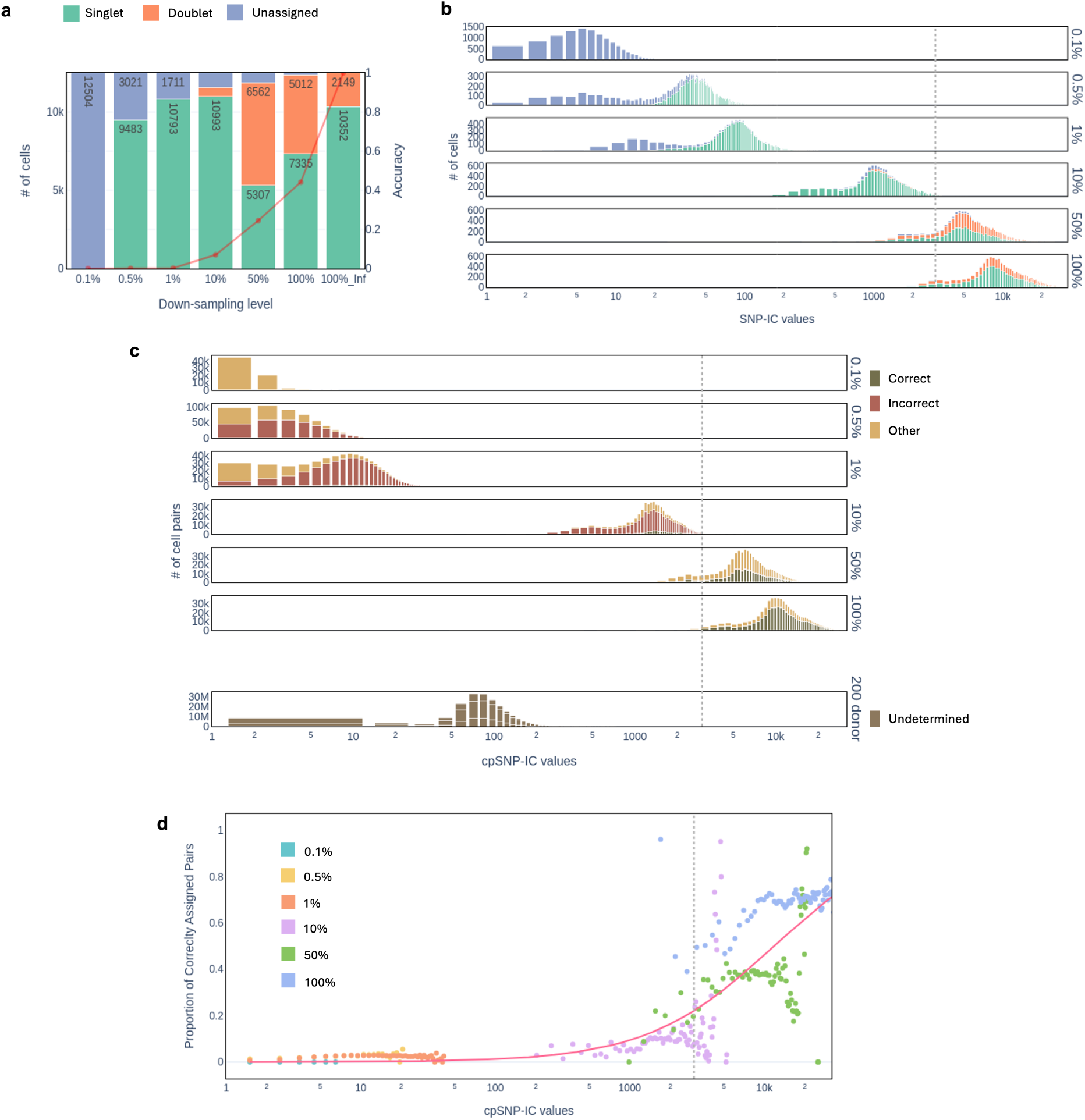
Demultiplex-ability analysis for genotype-free algorithms. (a) Genotype-free Vireo classification results (singlet, doublet, unassigned) in down-sampled datasets (0.1% - 100%) using the 104-donor cell village dataset. The demultiplexing accuracy is shown as a solid red line. The number of singlets peaks at 10%, although the accuracy increases monotonically. (b) Histogram of SNP-IC values for each down-sampled level. Different classification categories are shown in distinct colors. The singlet ratio does not increase monotonically with SNP-IC. The dashed line indicates cpSNP-IC = 50. Top: Histogram of cpSNP-IC values for each down-sampled level (top - 0.1%; last to bottom - 100% of reads). Different classification categories (correctly assigned, incorrectly assigned, others) are shown in different colors. The dashed line indicates cpSNP-IC = 3, 000, below which the classification are mostly incorrect or others. Bottom: The histogram of cpSNP-IC for 200-donor iPSC dataset. Their cpSNP-IC are far below 3,000, suggesting that the genotype-free Vireo fails to correctly cluster cells. Proportion of correctly assigned cell pairs versus cpSNP-IC for each down-sampled level in the 104-donor cell village dataset. The proportion begins to increase around cpSNP-IC = 3, 000.

Analyzing these effects at the single-cell level is more challenging for genotype-free methods than with-genotype methods, since the genotype-free methods group cells by assigning each cell a cluster index instead of a donor ID: In other words, distinguishing true singlets from false positives on a per-cell basis is difficult for genotype-free methods, especially when sequencing depth is low. Moreover, the algorithms behind genotype-free methods compare cells directly to other cells, so previously defined SNP-IC itself is not crucial to the degree to which a dataset can be demultiplexed. For these reasons, we introduce a quantity that, for each pair of cells, assesses how distinguishable the two cells are based on their shared SNPs. This quantity, cell-pair SNP-IC (cpSNP-IC), is computed by summing, over all SNPs they share, the smallest of the two UMI counts for each SNP (Fig. 1c). In other words, each SNP contributes according to the weaker signal observed between the two cells.

To study how cpSNP-IC affects demultiplexing performance, we start by generating all cell pairs, where both cells were singlets and belonged to the same donor as determined using the with-genotype algorithm with the original dataset (100%). We calculated the cpSNP-IC for each pair at each down-sampled level of the genotype-free results and generated the corresponding histograms, shown in Fig.4c. The histograms are stratified according to the results of genotype-free methods into three categories, visualized using different colors: *correct* if both cells are identified as singlets and assigned to the same donor; *incorrect* if both cells are identified as singlets but assigned to different donors; and *other* for all remaining cases, including when one or both cells are unassigned and/or identified as doublets). As shown in Fig.4c, cell pairs with low cpSNP-IC values are classified as either incorrect or other pairs. Correct pairs begin to emerge when cpSNP-IC exceeds 1000, which is much higher than the threshold we identified for SNP-IC. Fig. 4e shows the proportion of correct pairs as a function of cpSNP-IC, consistent with the trend observed in Fig. 4c. From this figure, we infer that a cpSNP-IC of approximately 3,000:

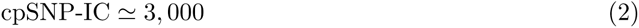

is necessary, at least, to reliably assign pair relationships. Interestingly, the figure also suggests that increasing cpSNP-IC, for example by increasing sequencing depth, could further improve de-multiplexing performance.

To validate our observation, we use our novel dataset of 200-donor human iPSC scRNAseq data. Since the sequencing depth is relatively lower than that of 104-donor dataset, not all cells meet criterion (2) (Fig.4c-bottom histogram). This explains the inconsistent results between different algorithms (Fig.S5-Rand index), as well as the observation that one donor appears to dominate the others. Overall, our criterion provides guidance on the minimum number of reads required based on the data prior to performing pooled experiments, which could help experimentalists design pooled scRNAseq experiments more effectively.

### 2.4 SNP microarray comparison

Our results show that with-genotype algorithms have a lower criterion of SNP-IC, suggesting their prioritized use when possible. But the use of these algorithms requires capturing the SNP genotype profile of donors. Since there are a number of different genotyping solutions currently available in the market, we briefly explore how sensitive are the results of the demultiplexation process to changes in the underlying SNP genotype space. For this, we selected four different microarray genotyping platforms: Infinium Global Diversity Array (GDA), Infinium Global Screening Array (GSA), Infinium OmniExpress Array (OEA) and the Infinium OmniExpressExome Array (OEE). More details on these genotyping arrays and the data processing are given in the *Methods section*. Looking at Fig.5a, we can clearly see the similarity between the mappings reached by all methods, expressed as the 4-way intersection of the singlets identified using each genotyping solution. This is also observed in the pairwise comparison between methods of Fig.5b. Here, the Jaccard index exceeds 98%, which indicates an almost complete overlap between the sets of singlets identified in each case. Moreover, the Rand index indicates that for singlets at the intersection between any two methods, the mapping is identical.

**Figure 5:**
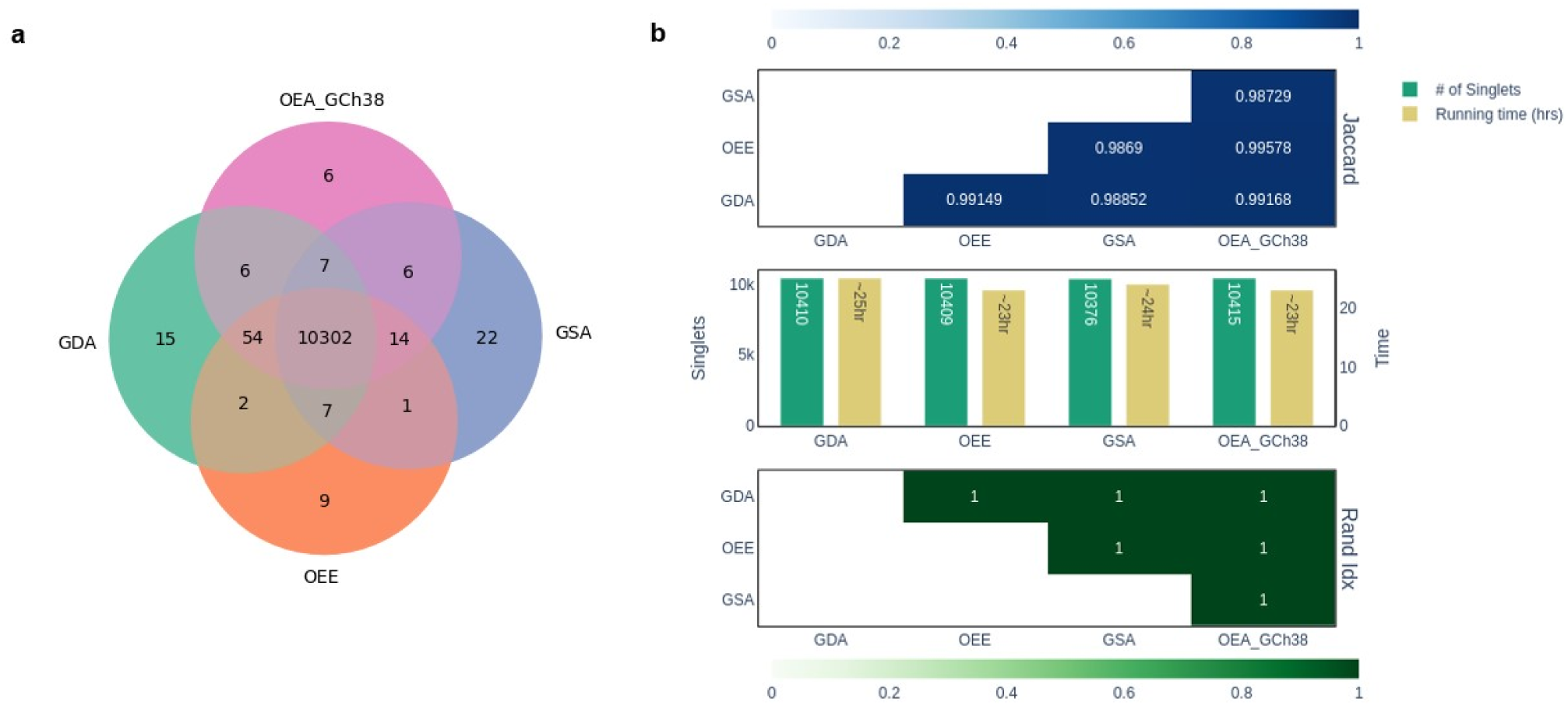
Comparison of with-genotype Vireo performance across different sets of SNPs. Various commercially available SNP microarrays were used for the 104-donor cell village dataset: Global Diversity Array (GDA), OmniExpressExome (OEE), Global Screening Array (GSA), and OmniExpressArray (OEA). (a) The overlap of cells identified as singlets for each microarray combination. (b) The Jaccard and Rand indices for pair-wise comparison of demultiplexing results indicate strong agreement between different platforms. In practice, we used the overlap between SNP sets derived from the 104-donor cell village dataset (obtained from whole genome sequencing) and the SNP list of each microarray. Please refer to the Methods section for details.

### 2.5 Framework and Package description

Finally, we propose *oddSNP* as a framework for the analysis of scRNAseq data and for optimizing the generation of pooled scRNAseq data, structured as shown in Fig.1d and Supplementary Fig.S6. The framework, implemented in Python and available as a stand-alone command line application, allows users to select and apply the methods discussed here to their individual scRNAseq datasets. Specifically, the software package provides functionality for down-sampling of scRNAseq data, specifying SNP microarrays, summarizing individual- and paired-cell genotype information (SNP-IC and cpSNP-IC) gathered from pileup, and plotting histogram summaries. It also performs simulations to estimate the proportion of cells that can be demultiplexed for each donor, using either the user-specified number of donors or a user-provided ratio vector. The implemented package serves as a wrapper for the core functions of *cellsnp-lite* that allow the generation of pileup results required for SNP-IC and cpSNP-IC calculation, as well as to *Vireo* routines for optional demultiplexation when pooled data are analyzed. (see Fig.S6 for details).

The source code for the implementation of *oddSNP* is freely available on GitHub [URL to be inserted]. Also, ready to use packages can be downloaded and installed through PyPi and bioconda.

## 3 Discussion

In this article, we propose a method to evaluate whether pooled scRNAseq data can be demultiplexed based on unpooled scRNAseq data from the same sample. This allows researchers to test, *a priori*, whether a demultiplexing algorithm will work under their experimental conditions. For this purpose, we focused on the information content of each cell (SNP-IC), which can be computed solely from unpooled scRNAseq data. We found that an SNP-IC of approximately 50 serves as a criterion for whether a cell can be demultiplexed using with-genotype algorithms. The general applicability of the method was tested using two different types of pooled scRNAseq, as shown in Figs 3d and 3e. For genotype-free algorithms, SNP-IC is not entirely informative as these methods work by comparing SNPs between cells. For this case, we introduce the information content shared between pairs of cells (cell-pair SNP-IC, or cpSNP-IC). We found that a cpSNP-IC of approximately 3000 is the minimum required to reliably assign pairwise relationships. To facilitate application of this test to any scRNAseq dataset, we provide oddSNP, a Python package that computes SNP-IC and cpSNP-IC for each cell barcode, available at [URL to be inserted].

To define SNP-IC for each cell barcode, we summed over the identified SNPs, weighted by the corresponding UMI counts. The underlying assumption is that each SNP contributes equally to the ability to correctly assign cell barcodes. While this is an approximation, a more precise measure would account for the specific alleles considered (e.g., allele frequency) as well as the composition of other donors in the pool. For the former factor, incorporating minor allele frequency into the SNP-IC calculation as a weight would be straightforward. Nevertheless, our criterion for SNP-IC appears to be relatively robust to both allele details and donor composition, as demonstrated by two independent datasets with mixtures of 24 or 104 donor (Fig.3d-e).

With-genotype algorithms exhibit relatively robust performance that does not depend on the specific factors we tested, such as the type of algorithm (Fig.2) and the number of pooled donors (Fig.S2). However, it is important to note that the sequencing depths in these comparisons are relatively high. At lower sequencing depths, the extent to which these parameters influence performance remains an open question and represents an interesting area for future study. Addressing this could help better determine optimal conditions for both experimental design and data analysis. The development of new algorithms for demultiplexing should also be benchmarked under low-depth sequencing conditions. In this context, the down-sampling strategies we employed (Fig.3 and Fig.4) would be useful.

We acknowledge the lack of a definitive ground truth throughout our demultiplexing analysis. However, by adopting a consensus approach (as in [17]) and employing statistical measures that provide insight into mapping similarity, we are able to establish a reliable background against which comparisons can be effectively made. This approach has allowed us to assess the validity of both with-genotype and genotype-free algorithms when applied to datasets containing a number of pooled donors far greater than the fewer than 10 donors, for which experimental validations have been reported so far [17].

We found execution time and memory load to remain as important issues, especially when looking at datasets that continue to increase in size. In this context, one important line of improvement relates to fine tuning the selection of reference variants used for pile-up processes; a component that lies at the base of both oddSNP and with-genotype algorithms in general.

Finally, it is crucial to note that the threshold for SNP-IC in with-genotype algorithms and cpSNP-IC in genotype-free algorithms differs by two orders of magnitude, suggesting the superiority of with-genotype algorithms when genotype information is available. This difference arises because cpSNP-IC compares two cell barcodes, both of which contain incomplete SNP profiles, whereas SNP-IC compares a cell barcode against a reference panel with a very rich SNP profile, including millions of SNPs. As a result, genotype-free algorithms often fall into a regime where the cpSNP-IC for each cell pair is not sufficiently large (*cf.*, Fig.4). However, our results suggest that this limitation could be mitigated by increasing sequencing depth, which is not the most expensive component of scRNAseq experiments.

## 4 Methods

### 4.1 Demultiplexing Methods

Polymorphism-based demultiplexing methods use SNP signatures of cells to group them and map them to their corresponding donors. With-genotype methods use specific SNP locations, where the donors genotypes are known, to generate these signatures; whilst genotype-free methods use SNP locations from across a whole population reference.

In this article, we use the following with-genotype methods for demultiplexing:

- **Demuxlet** is a method that uses maximum likelihood to determine the most likely donor for each droplet in a pooled dataset using a mixture model. The method usually involves performing a two-step process: a pileup step used to identify the number of reads at each SNP location, followed by a demultiplexing step [18].
- **Dropulation**, like Demuxlet, also uses a maximum likelihood approach that uses pre-existing whole genome sequence or SNP data to assign cells to donors based on the likelihood of the observed combination of transcribed alleles to have been generated from the donor genome sequence [13].
- **Demuxalot** authors model the demultiplexing inference problem based on a generative process of the aligned reads, and use this to develop a Bayesian framework to infer the genotype of each initial cell barcode [20].

We will also use the following list of genotype-free methods:

- **Freemuxlet** uses an unsupervised clustering strategy, specifically a modified Expectation-Maximization algorithm, to assign cell barcodes to clusters in an iterative cycle, defining singlets as cells assigned to a single cluster, and multiplets as cells that cannot be assigned unequivocally to a single cluster [21].
- **ScSplit** uses the list of variants called from the original sequencing data itself. Using the allelic counts computed based on these variants, it assigns samples to clusters using an Expectation-Maximization framework [23]. The process can also be improved by filtering the called variants with genomic information coming from general populations.
- **Souporcell** starts by remapping and calling variants on the original reads, to then count the cell alleles. Souporcell then uses the computed cell allele support counts to cluster cells using a sparse mixture model clustering [22].

Finally, we also consider a method that can perform the demultiplexing process in a genotype-free and with-genotype setting:

- **Vireo** uses common genetic variants to reconstruct the partial genotypic state of individuals in the donor pool and then uses a variational Bayesian inference to assign each cell to one of these individuals [19]. Vireo also allows for the use of partial or complete genotypic information about the donors of the pool.

### 4.2 Pooled scRNAseq Input Dataset

#### 4.2.1 104-donor cell village dataset

We used 104-donor *cell village* dataset from Wells *et al.*, consisting of human iPSCs [13]. For the 104-donor cell village, the authors first expanded the iPSC line from each donor separately for one week, followed by dissociation into single-cell suspensions. The cells were then counted using an automated handheld cell counter. To create a uniformly mixed village, equal numbers of cells from each of the 104 donors were combined and plated together. The pooled culture was grown and stabilized for five days. Finally, the mixed cell population was harvested and loaded onto the 10x Chromium platform for droplet encapsulation and single-cell RNA sequencing.

The authors have made publicly available scRNAseq datasets that combine cells from different donors, together with genotype information, in the form of SNP profiles. The original data included scRNAseq from 15 individual cell capture experiments. Here we only use the data corresponding to the cell capture experiment *A10*. The downloaded data are over 218 GB in size, including 104 donors and a total of more than 3.6 million cells. The authors also provide a cell-to-donor mapping for a subset of 9,692 cells.

Although cell-to-donor mapping is originally provided for 9,962 cells, it is not clear if or how this list of cell barcodes was selected from the original list of scRNAseq reads. To generate a similar, filtered list of target cells to use for demultiplexing, we used a simple quality assessment approach, whereby we first filtered the original reads to keep only cells that had a UMI count over 25,000. Then, we further filtered cells to include only those that have a gene count (number of expressed genes) between 4,000 and 12,000. The final list of selected barcodes includes 10,169 cells, from which 8,894 are also found in the original list. Supplementary Fig. S7 shows a UMI count and gene count comparison between the selected target cells and the original ones. This dataset includes genotype information for all donors.

#### 4.2.2 Human liver organoid dataset

In addition, we will use a scRNAseq dataset of iPSC derived HLO that corresponds to data reported in [16]. This dataset includes pooled information from 24 donors for a total of 2.1 million cells (with a subset of 14,504 target cells). Although this dataset does not include a cell-to-donor mapping for target cells, it does include genotype information for all donors.

Note that for the generation of panels (b) and (c) in Fig.3, we considered only cells classified as singlets in the original dataset (100% of reads), while for panel (d), we also included cells classified as either singlets or unassigned in the original dataset.

#### 4.2.3 Generation of 200-donor human iPSC dataset

For the final dataset, human iPSC was derived from primary human fibroblast cells obtained from 200 donors (Cell Systems, Kirkland, WA, USA). Primary human fibroblast cells were cultured and expanded according to the manufacturer’s instructions. The expanded fibroblast cells were reprogrammed to iPSC as described previously [27]. The established iPSC was harvested within five passages and used for the preparation of RNA sequencing. For the construction of single-cell RNA-Seq library, Chromium Next GEM Single Cell 3’ Library and Gel Bead Kit v3.1 (10x Genomics) was used according to the manufacturer’s instructions. Libraries were sequenced on a DNBSEQ-G400 (MGI) at a read length of 28x100 bp paired-end to yield a minimum of 20,000 reads per cell for gene expression.

This dataset (add publication link) includes a total of more than 2.4 million cells (and a subset of 20,880 target cells). This dataset does not include information on the genotype of donors.

A summary of all scRNAseq input datasets is given in Table 1.

**Table 1:** Dataset summary information.

| Dataset | Nº donors | Nº Raw Cells | Nº Target Cells | Size |
| --- | --- | --- | --- | --- |
| 104 cell-village | 104 | ~3.6M | ~12K | ~218GB |
| 24 HLO | 24 | ~2.1M | ~14K | ~26GB |
| 200 iPSC | 200 | ~2.8M | ~20K | ~50GB |

### 4.3 Genetic variation (SNP) reference

Natural genetic variation demultiplexing methods are based on the idea that variation patterns are kept constant among cells that belong to the same donor. In order to identify the patterns, we first need to define the variation sites within the genome, that is, we need to define the list of reference SNPs that we will use.

Here, we use a filtered selection of variation sites generated on data from the 1000 Genome project. Genetic variation sites were generated by aligning the 1000 Genome project sequence data with the GRCh38 reference genome and performing variant calling directly against it. The set is then restricted to single nucleotide variants and indels. This reference includes more than 78 million variants. Data are publicly available for download from the International Genome Sample Resource (IGSR - https://www.internationalgenome.org/)[28].

### 4.4 Demultiplexing algorithms execution

We apply all demultiplexing algorithms to the 104-donor cell-village dataset (*cell capture experiment A10* [13]). All algorithms were executed using a single configuration of 15 cores, for algorithms that include such parameter in their execution commands, and two upper limits of 400 GB and 750 GB of available RAM (See Methods Section 4.8).

Each demultiplexing algorithm except Dropulation was run on the selected subset of 10,169 cell barcodes. For Dropulation, we use the demultiplexing results of the original article [13]. The same list of reference variants was used for all algorithms (see Methods Section 4.3), complemented by genotype information from the original studies [12, 13] for the case of with-genotype algorithms. Supplementary Fig. 2 shows a comparison of the demultiplexing results obtained using different algorithms (panels a and c), along with a summary of the running time required for each algorithm (panel b).

With the exception of Demuxlet and Freemuxlet, all the methods tested were able to complete execution (Running Demuxlet encountered memory issues when using a 750 GB upper limit of available RAM, whilst Freemuxlet execution was forcefully stopped after 10 days of execution). The number of identified singlets varies from approximately 6,600 for the uninformed version of Vireo, to nearly 8,900 in the informed versions of Vireo and Demuxalot, as shown by the bar graph in Fig. 2.

### 4.5 Additional Genotyping Sources

With-genotype demultiplexation methods can only be used when donor genotype information is available. Genotype information for the 104-donor cell village dataset is made available in the form of a list of more than 31 million variants generated based on calling the original sequences against the GRCh38 reference genome [13]. Genotype information is also available for the 24-donor HLO dataset. In this case, the list of informed variants corresponds to those measured by [16] using TaqMan SNP Genotyping Assays (Thermo Fisher Scientific Inc.). This domain includes almost 950,000 variation sites. Of these, 838,913 variation sites are found in the intersection with the genetic variation reference.

We also generated a series of alternative genotype information files for the 104-donor cell village dataset, and they constitute the base for the SNP microarray comparison section. Each alternative genotype is generated using the intersection of the variants originally provided by the authors in [13] and one of the following microarray genotyping solutions: Infinium Global Diversity array (GDA), Infinium Global Screening array (GSA), Infinium Omniexpress (OEA), and Infinium Omniexpress Exome (OEE). Table 2 provides a detailed summary of the microarray version used and the number of variants included in each of them.

**Table 2.**

| Array | Version | Original № of markers | Intersected variants |
| --- | --- | --- | --- |
| GDA | 8v1-0_D2 | ~1.8 million | ~1.6 million |
| GSA | 24v3-0_A2 | ~650,000 | ~517,000 |
| OEA | 24v1-4_A1 | ~720,000 | ~660,000 |
| OEE | 8v1-6_A2 | ~960,000 | ~720,000 |

For with-genotype Vireo, we use the list of reference variants common to all of our experiments.

### 4.6 SNP information content (SNP-IC) and cell-pair SNP-IC

Given a specific variation site in the genotype reference file, the software package cellSNP-lite is able to calculate the number of times the reference, alternative, and other alleles appear at that site in a scRNAseq dataset. These pileup counts for each site (filtered to the variants common to donor genotype, when available) are stored in a series of matrix files, ready to be used by Vireo for demultiplexing.

To calculate the SNP-IC for any given cell *i*, we take the counts for each SNP stored in the pileup files and calculate their sum, *i.e.* their total UMI count, then for all SNPs reported for a single cell, we again sum the counts for each SNP to obtain the final SNP-IC value:

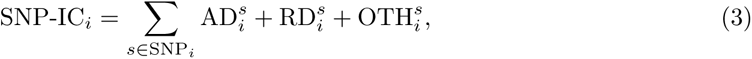

where AD*_i_^s^* is the number of times that SNP *s* was found to have the alternative allele in the current cell *i*, RD*_i_^s^* is the number of times the reference allele was found, and OTH*_i_^s^* is the number of times other allele was found. The set SNP*_i_* represents all SNPs identified in the given cell *i*. Calculating the SNP-IC for each cell in a given scRNAseq dataset can be done by using command snpic in oddSNP.

Similarly, the cell-pair SNP-IC between any given cell pair (*i* and *j*) is calculated as the sum, over all shared SNPs, of the smaller UMI count between the two cells for each SNP:

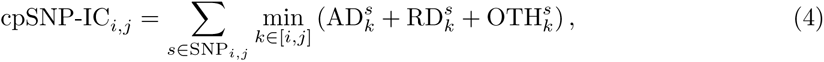

where the set SNP*_i,j_* represents the intersection of SNPs found on cells *i* and *j* (i.e. SNP*_i,j_* = SNP*_i_* ∩ SNP*_j_*).

### 4.7 Evaluation Metrics

Taking as a base the cell-to-donor mapping provided by the authors, we will use statistics that measure the degree of mutual agreement between this mapping and the mappings generated at the different stages of analysis to evaluate the performance of the different algorithms. Specifically, we use the *Rand index* and the *Jaccard index*.

#### 4.7.1 Rand Index

Named after William M. Rand, the rand index is a measure of similarity between two different data clustering results [26]. For two different clusterings of the same set of points, the Rand index indicates the degree of similarity between them by taking pairs of elements and checking if they are placed together (or separated) in the same way in both clusterings. The Rand index takes a value between 0 and 1, with 0 meaning that there is no agreement in the clustering of any pair of points, and 1 meaning that the clusterings are the same.

#### 4.7.2 Jaccard Index

Also known as the *Jaccard similarity coefficient*, is a statistic used to measure the similarity between two sets that is calculated by taking the size of the intersection of the two sets and dividing it by their union. Like the Rand index, the Jaccard index also takes a value between 0 and 1, with 0 meaning that the sets are completely different, and 1 meaning that the sets are the same [25].

### 4.8 Specification of computational resources

Execution runs with a memory limit of 400 GB were carried out using a 64-bit Intel Xeon Gold 6238R CPU @2.20GHz machine with 112 processing units, 500 GB of RAM and an NVIDIA RTX A6000 graphics card; whilst execution runs with an upper limit of 750 GB of memory were carried out using a similar 64-bit Intel Xeon Gold 6238R CPU @2.20GHz machine with 112 processing units and 1 TB of available RAM.

## 5 Data Availability

The two datasets used in this study have already been published in [12] (104-donor cell-village dataset) and [16] (24-donor liver organoid pooled dataset). Our in-house 200-donor iPSC dataset will be uploaded to GEO database (accession number to be added) upon publication.

Source code for the implementation of *oddSNP* is freely available on GitHub [URL to be inserted]. Also, ready to use packages can be downloaded and installed through PyPi and bioconda.

## 6 Acknowledgments

We would like to thank Michael F. Wells for providing the details of the preprocessing pipeline for the 104-donor cell village dataset. We acknowledge the NGS core facility at the Research Institute for Microbial Diseases of The University of Osaka for the sequencing and data analysis.

## 9 Supplementary Information

**Figure S1:**
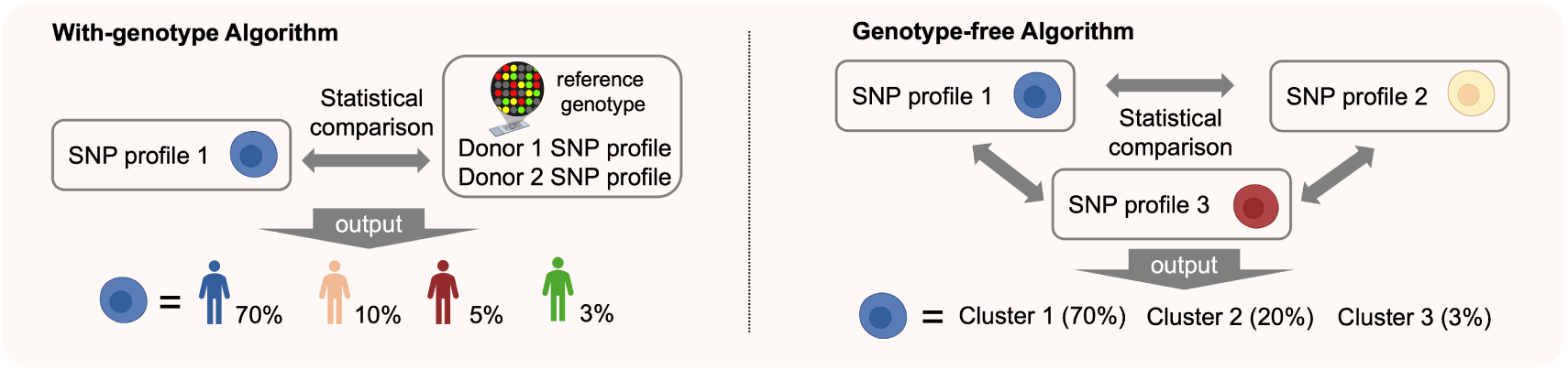
With-genotype algorithms and genotype-free algorithms. In with-genotype strategies (left), the SNP profile of each cell is compared with reference genotypes obtained from donors in advance. In contrast, genotype-free strategies (right) compare SNP profiles between cells to cluster them and infer which cells originate from the same donors.

**Figure S2:**
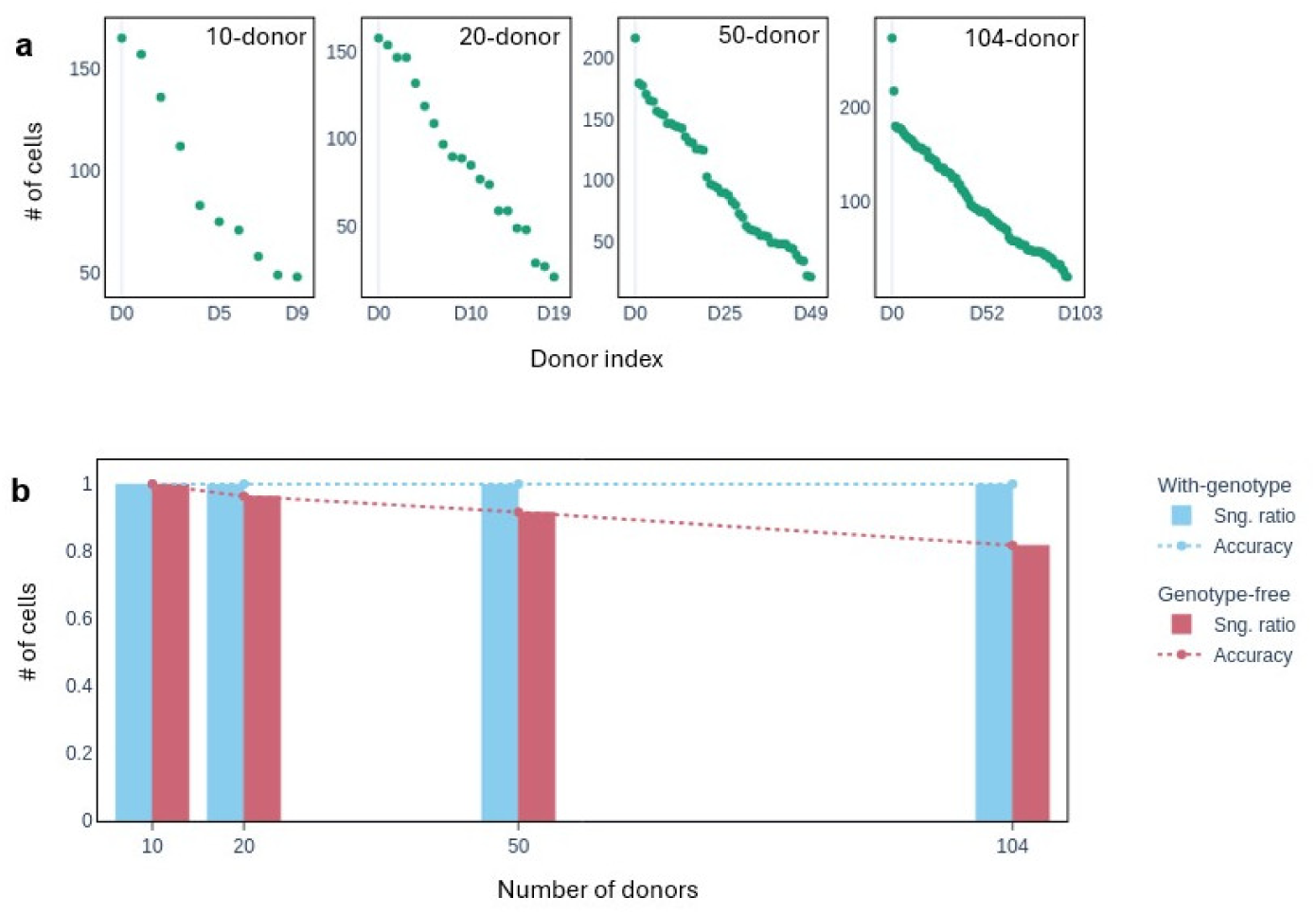
To demonstrate the impact of the number of pooled donors, we generated a series of down-sampled datasets from the original scRNAseq data by randomly selecting 10, 20, and 50 donors. In doing so, we preserved the linearly decreasing trend in the number of cells per donor, as shown in the panel (a), where the numbers of cells per donor across 4 donor-downsampled datasets (10, 20, 50, and 104 (original)) are shown. Donor indices are reordered in descending order based on the number of cells. This ensured that SNP information content remained the same. We then applied both with-genotype and genotype-free versions of Vireo to demultiplex each dataset and compared the resulting assignments those obtained using all original donors. The panel (b) shows how the demultiplexing results change when using the with-genotype and genotype-free versions of Vireo across the down-sampled datasets. The proportion of cells classified as singlets (bar graphs) and the demultiplexing accuracy (dotted lines), defined as the product of the Jaccard and Rand indices. Both quantities are shown for with-genotype (blue) and genotype-free (red) algorithms as a function of the number of donors (using the 4 donor-downsampled dataset). As expected, the with-genotype method does not change the demultiplexing performance as the number of donors decreases, since the SNP information content remains similar. By contrast, the genotype-free method shows improved accuracy when the number of donors decreases. This suggests that, in addition to SNP information, the composition of the donor also influences demultiplexing performance.

**Figure S3:**
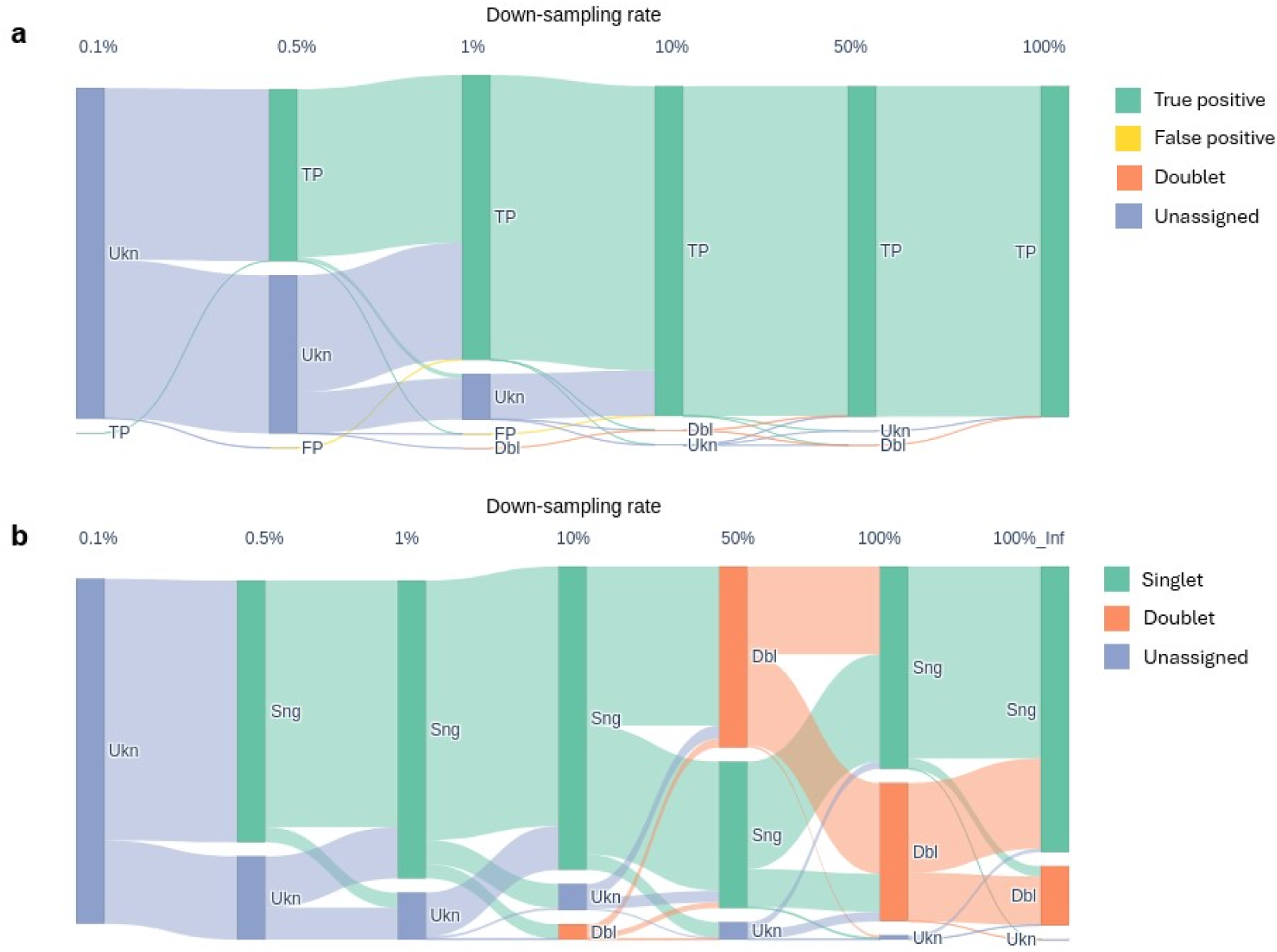
Changes in cell classification between consecutive down-sampled levels are visualized using a Sankey diagram. (a) Classification results from with-genotype version of Vireo. (b) Classification results from genotype-free version of Vireo.

**Figure S4:**
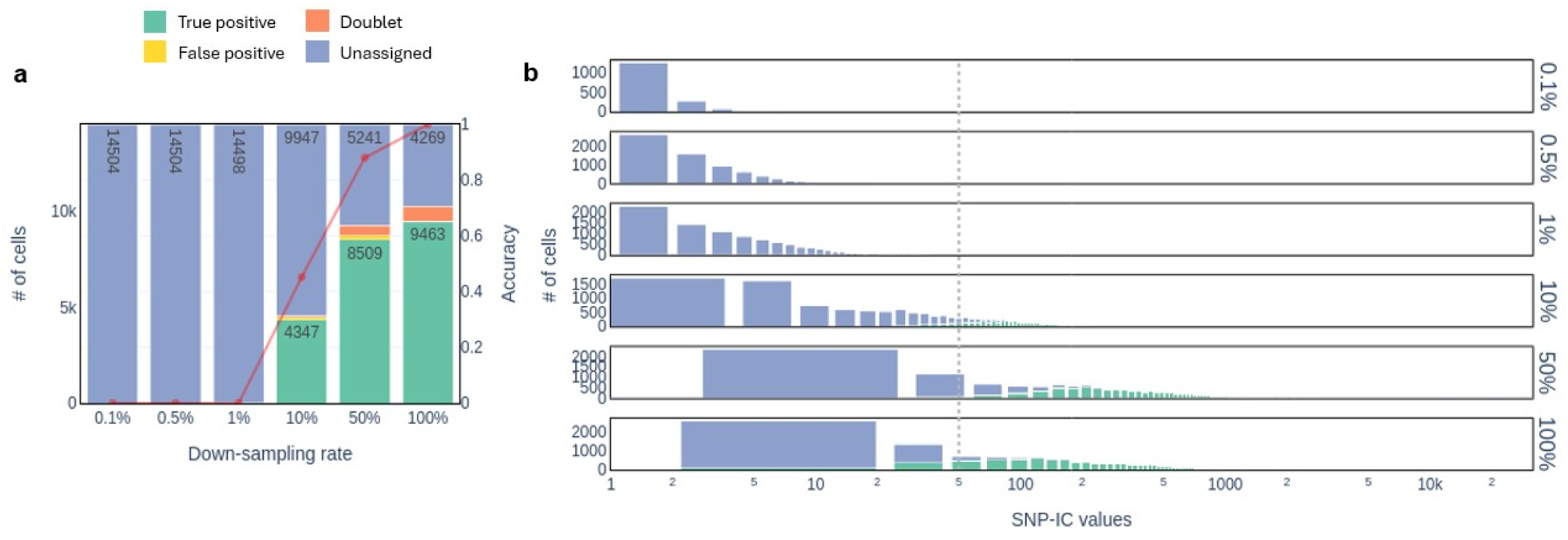
Results for the 24-donor HLO dataset. (a) Cell-type observed for with-genotype, Vireo demultiplexation at each down-sampled level of reads. (b) Histogram of SNP-IC for each down-sampled level.

**Figure S5:**
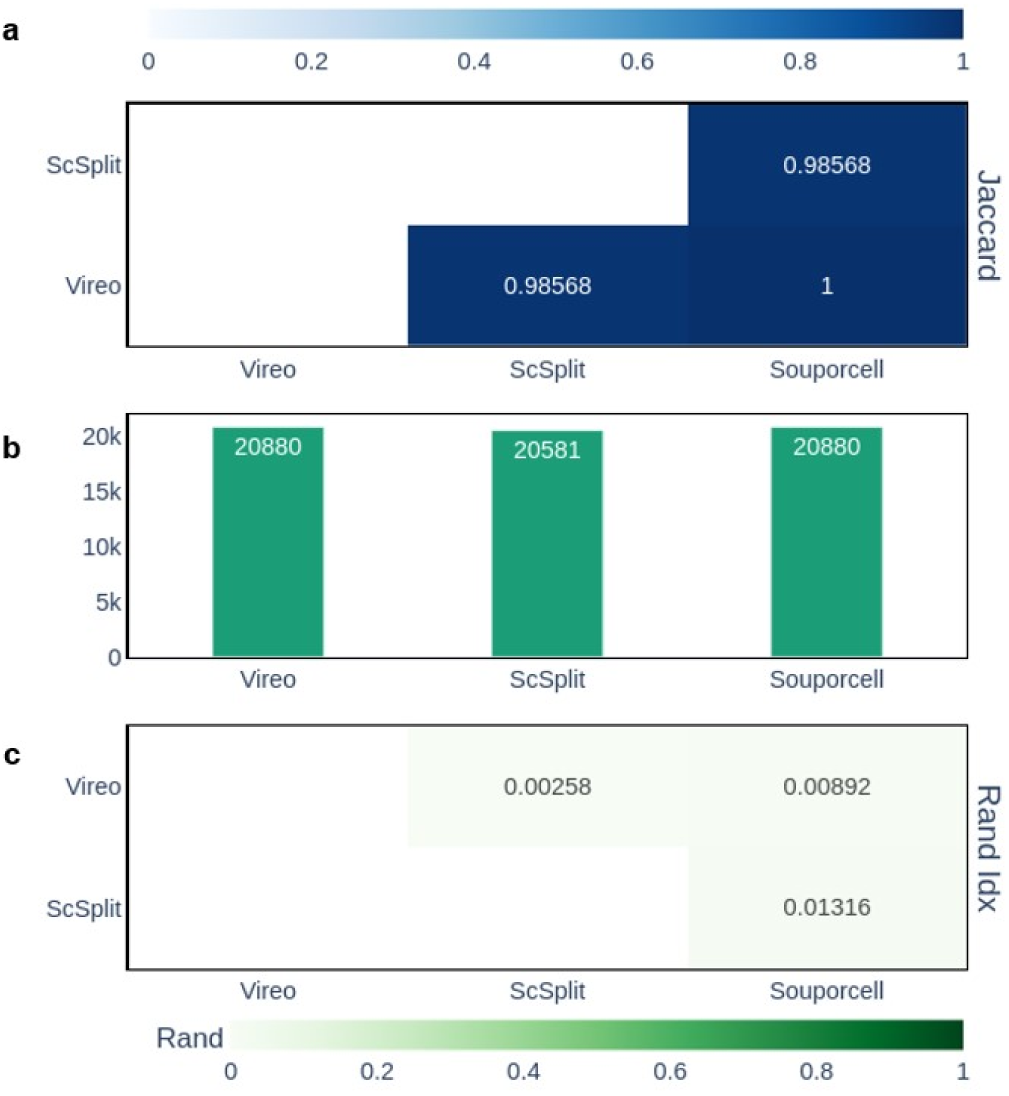
Genotype-free demultiplexing results obtained for the in-house 200-donor HLO dataset.

**Figure S6:**
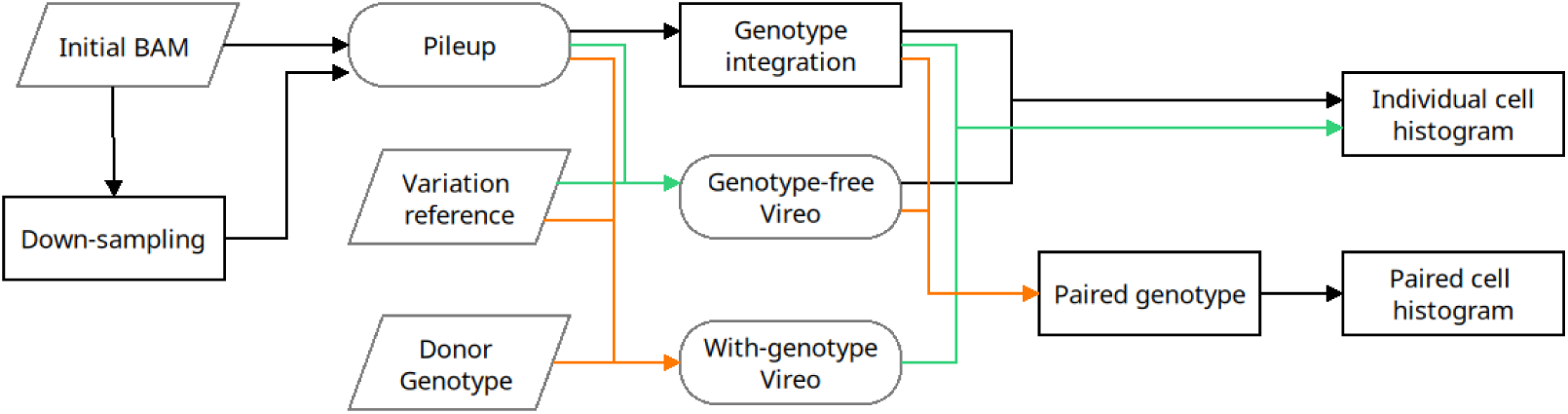
Pipeline structure for oddSNP. Boxes are used to indicate routines implemented specifically for oddSNP; rounded boxes indicate integration with previously published tools.

**Figure S7:**
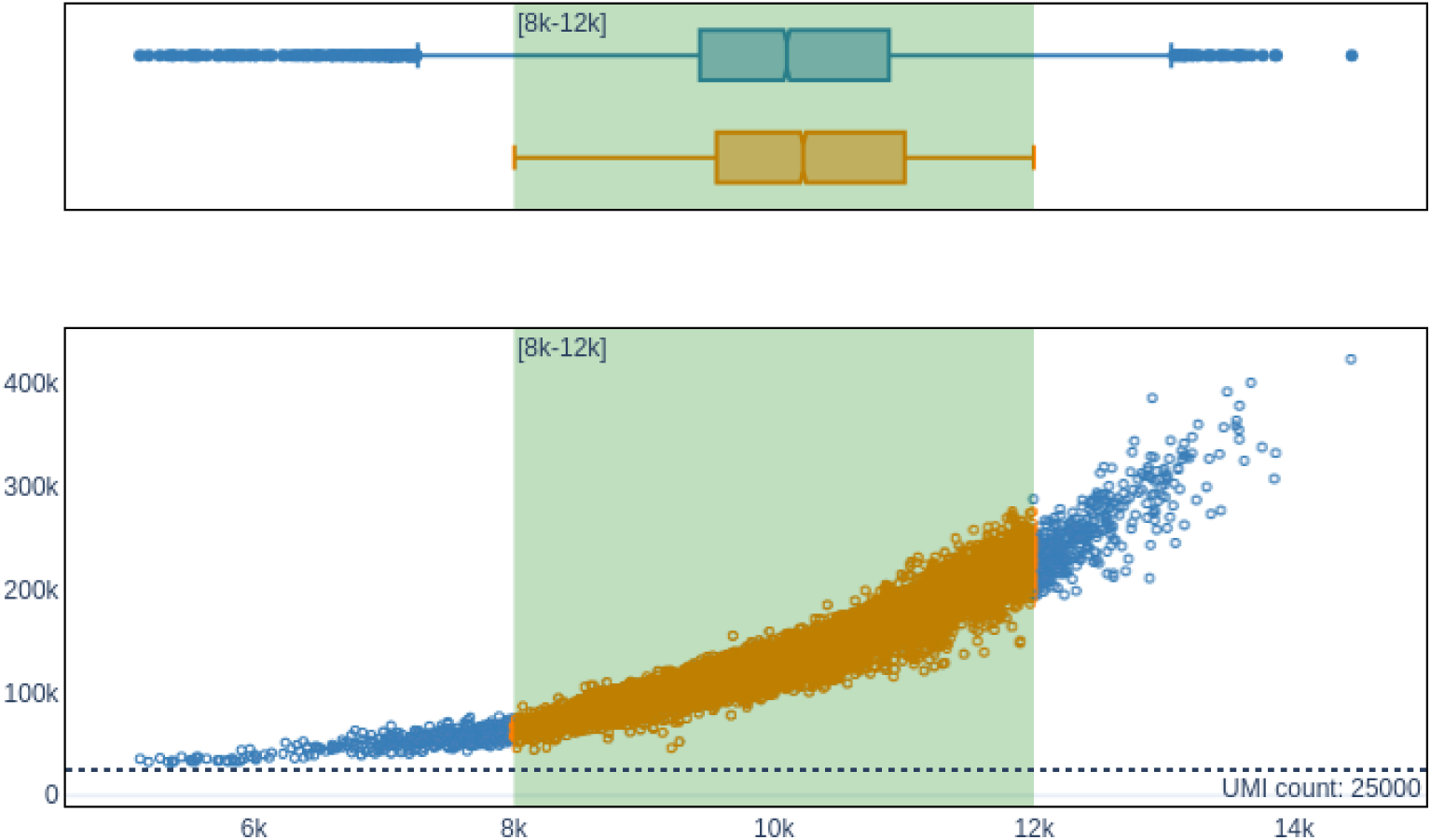
Cells from 104-cell village dataset selected for demultiplexation. Originally demultiplexed cells are shown in blue, and selected cells in orange. Only cells that express more than 4000 genes are included for plotting.

